# Baktfold: Sensitive protein functional annotation across the microbial tree of life using structural information

**DOI:** 10.64898/2026.03.31.715528

**Authors:** George Bouras, Sung won Lim, Lindsay Durr, Sarah Vreugde, Alexander Goesmann, Robert A. Edwards, Oliver Schwengers

## Abstract

The functional annotation of protein sequences has undergone tremendous progress over recent years, but still too-many protein sequences remain as so-called hypothetical proteins after applying state-of-the-art genome annotation software pipelines. Here, we introduce Baktfold, a new command line software tool for the ultra-sensitive but taxon-independent fast annotation of protein sequences across the microbial tree of life. Baktfold conducts sequential protein structure-based searches against four complementary structure databases. Protein sequences are transformed into Foldseek 3Di tokens via the ProstT5 protein language model and subsequently searched against structure databases via Foldseek. All results are exported in GFF3 and INSDC-compliant flat files as well as comprehensive JSON files facilitating automated downstream analysis 100% interoperable with the popular bacterial annotation tool Bakta. We compared Baktfold’s performance in terms of wallclock runtime and functional annotation of hypothetical proteins from various sources including bacterial and archaeal isolates, plasmids, metagenomic-assembled genomes and micro-eukaryotes. When benchmarked on over three hundred thousand species representatives across the prokaryotic tree of life, Baktfold’s median overall bacterial genome annotation rate is 87.8% compared to 72.9% with Bakta, while Baktfold’s median bacterial annotation rate of remaining hypothetical proteins is 50.1% (n=290258). For archaea, Baktfold’s overall median annotation rate is 71.5% compared to Prokka’s 35.8%, with a median archaeal annotation rate of hypothetical proteins of 68.0% (n=14058), making Baktfold the most sensitive automated archaeal annotation method by far. Baktfold is implemented in Python 3 and runs on MacOS and Linux systems. It is freely available under a MIT license at https://github.com/gbouras13/baktfold.

**Data Summary:**

1. Baktfold was developed in Python as a command line application for Linux and MacOS
2. The complete source code and documentation are available on GitHub under an MIT license: https://github.com/gbouras13/baktfold
3. The Baktfold database is hosted at Zenodo (https://zenodo.org/records/17347516) mirrored on HuggingFace (https://huggingface.co/datasets/gbouras13/baktfold-db)
4. Baktfold is available via bioconda (https://anaconda.org/bioconda/baktfold) and PyPI (https://pypi.org/project/baktfold/)
5. Baktfold can also be run without local installation using Google Colab at https://colab.research.google.com/github/gbouras13/baktfold/blob/main/run_baktfold. ipynb
6. All supplementary code, data and files required to reproduce the results of this manuscript are available at https://github.com/gbouras13/baktfold-analysis (code and small data) and https://zenodo.org/records/19333697 (large data)

## Introduction

As sequencing data from both cultured isolates[1] and metagenomic sources[2]continues to expand, our ability to characterise the functional capacity of microbial life is increasingly challenged; rapid and automated genome annotation has therefore become essential. Widely used command-line tools such as Bakta[3], Prokka[4] and DFAST[5] enable fast and consistent annotation of bacterial and archaeal genomes. These tools primarily rely on protein sequence homology, leveraging approaches such as sequence hashing (as implemented in Bakta[3]), sequence alignment (e.g. BLAST[6], DIAMOND[7], MMseqs2[8]), and profile hidden Markov models (e.g. HMMER[9, 10], HHblits[11]). Tools such as eggNOG-mapper[12] further incorporate orthology-based inference to improve protein annotation across diverse taxa, albeit with greater computational and database requirements compared with more nimble tools like Prokka and Bakta.

Despite these advances, a substantial fraction of microbial proteins remain functionally uncharacterised, with estimates suggesting that approximately 30% of bacterial proteins lack functional annotation[13, 14]. As one moves further towards the frontier of microbial protein function prediction, this challenge is even more pronounced, with archaeal [15–17] and microbial eukaryotic[18] proteins annotation proving particularly difficult.

Recent advances in protein structure prediction, catalysed by the success of AlphaFold2[19], have enabled accurate structure prediction directly from sequence. Because protein structure is more conserved than sequence and more closely linked to function[20], structural information provides a powerful avenue for annotating proteins beyond the limits of sequence homology. The availability of large-scale structure databases, including AlphaFold Protein Structure Database (‘AFDB’)[21, 22] and Protein Data Bank (‘PDB’)[23], combined with fast structural search tools such as Foldseek[24], now enables detection of homology deep within and beyond the “twilight zone” of sequence identity (defined as 20-35% sequence identity)[25].

However, protein structural information is not yet routinely integrated into automated genome-scale or metagenome-scale annotation pipelines. Existing structure prediction tools such as ESMFold[26] and ColabFold[27], while revolutionary, highly accurate, accessible and open-source, remain computationally demanding when applied at genome or metagenome scale (including large scale GPU and database storage requirements), limiting their use in high-throughput annotation workflows.

Protein language models (pLMs), such as ProtT5[28] and ESM-2[26] have emerged as pillars upon which protein function prediction tools may be built. pLMs are transformer-based models trained on large protein sequence corpora, learning representations that capture structural[29] and functional properties[30]. They therefore enable annotation beyond traditional homology-based transfer approaches[31], particularly for divergent or previously uncharacterised proteins.

A growing number of tools leverage pLM embeddings for functional annotation, including Empathi[32], FANTASIA[33], GAIA[34], and DeepFRI[35]. For example, FANTASIA transfers Gene Ontology[36] (GO) annotations using embedding similarity, while GAIA performs nearest-neighbour annotation in embedding space. DeepFRI combines pLM-derived features with structural information using graph convolutional networks to predict GO terms and localise functional sites. Despite their strong performance, most existing methods are limited to predicting specific annotation types—most commonly GO terms—and often produce probabilistic or embedding-based outputs that are less directly interpretable than traditional alignment-based approaches[37].

Here, we present Baktfold, our tool for rapid and standardised protein annotation across the microbial tree of life that integrates structural information at genome and metagenome scales. Baktfold uses the ProstT5 protein language model[28] to predict structural representations in the form of Foldseek[24] 3Di sequences, which are combined with amino acid sequences and searched sequentially using Foldseek against multiple databases including AFDB clusters[38], Swiss-Prot[39], PDB[23], and CATH[40], with the user able to provide custom databases in addition.

We demonstrate that Baktfold complements and extends traditional sequence-based homology annotation approaches whilst providing interpretable alignment-based outputs. Across diverse datasets initially focusing on bacteria, but also spanning, archaea, plasmids, and microbial eukaryotes, Baktfold substantially improves functional annotation rates compared to sequence-based homology annotation. For example, the median annotation rate across GlobDB[41] bacterial species representatives increases to 87.8% compared to 72.9% using Bakta. The difference is even more pronounced for archaeal species representatives (71.5% compared to 35.8% using Prokka).

Baktfold supports both annotated genome (Bakta’s JSON and GenBank format) and protein FASTA inputs and produces standardised outputs compatible with downstream analysis and database submission, facilitating large-scale integration of structurally informed annotations into public databases.

## Materials and Methods

Across this manuscript, ‘functional annotation rate’ means the proportion of coding sequences (CDS) with a functional label not equivalent to “hypothetical protein” (in the context of Bakta, Prokka and Baktfold). This is strict, so even if the CDS has a database hit, if that hit has no functional label (and so is called hypothetical protein), this is not counted as a functional annotation. ‘Overall annotation rate’ as used in the eukaryotic genome analysis section, refers to all CDS with a target database hit for Baktfold and eggNOG-mapper, regardless of if the target database possesses a functional label.

All Baktfold benchmark runs, unless otherwise stated, were conducted using a high-performance computing (HPC) node on Setonix at the Pawsey Supercomputing Centre with AMD MI250x GPUs (one GPU was used with Baktfold) along with 8 cores (16 threads) and 32 GBs CPU memory of a 2.45GHz AMD EPYC 7763 “Milan” 64-Core CPU Node. Baktfold was run using a container with GPU utlising ROCM-compatible Pytorch v2.7.1 available at https://quay.io/repository/gbouras13/baktfold.

Two analyses were not run on Setonix related to Figure 4 i.e. the runtime wallclock analysis and the comparison between Baktfold with ProstT5[28], ColabFold/AlphaFold2[19, 27] and ESMFold[26] structures. These analyses were conducted on a HPC node on Phoenix HPC at Adelaide University (Australia) using a single NVIDIA A100-40G GPU, along with 32-threads CPU and 128 GB RAM of 2× Intel Xeon Platinum 3460Y CPU system with 36 cores @ 2.4GHz.

### Baktfold Database Construction

The Swiss-Prot (using AlphaFold Database v6 release)[21, 39], CATH[40] and PDB[23] databases were downloaded on 9 October 2025 using the ‘foldseek databases’ commands with Foldseek[24] version commit v8979d2 and included as is into Baktfold’s database. For the AlphaFold Database clusters, these were redone using the v6 release update following the same procedure as the original AFDB clusters manuscript[38] with Foldseek version v8979d2. Briefly, we took the AFDB50 cluster representatives downloaded using ‘foldseek databases’ (MMseqs2[8] clusters using 50% minimum sequence identity with 90% minimum sequence overlap). We then clustered these using ‘foldseek cluster’ with E=0.01 and 90% minimum structure overlap. From here, we kept one representative for each non-singleton cluster that was not indicated as a protein fragment. Due to the increase in size of the AFDB in v6[21], this yielded 3,085,778 representative proteins that were included in the Baktfold databases.

### GlobDB Dataset Analysis

All GlobDB[41] genomes were downloaded in FASTA format from https://globdb.org/downloads and annotated with Bakta v1.11.4[3]. For every genome that successfully completed annotation with Bakta (n=305,825/306,259), we then ran Baktfold v0.1.0 using the Bakta JSON output with default parameters. We also downloaded the publicly available Prokka[4] v1.14.6 annotations from https://opendata.eawag.ch/dataset/globdbr226-genbank-files for all genomes. This was specifically for comparison with Baktfold for the 14,812 archaea genomes, in order to use Prokka as a fairer annotation baseline given Bakta was not designed for archaeal genome annotation.

### IMG/PR Plasmid Protein Analysis

All plasmid proteins from IMG/PR[42] were downloaded from https://genome.jgi.doe.gov/portal/IMG_PR/IMG_PR.download.html on 16 March 2026. All proteins were then deduplicated using Seqkit2’s ‘seqkit rmdup’ command[43], followed by removal of a small number of proteins containing internal stop codons, yielding 8,809,078 proteins. Bakta v1.11.4[3] was then run using ‘bakta_proteins’ command, while Baktfold v0.1.0 was run using the ‘baktfold proteins’ command.

### Gene Ontology (GO) Term Analysis

For each GlobDB CDS that had a Baktfold hit to Baktfold’s Swiss-Prot database, we mapped the Swiss-Prot accession to the constituent GO terms using the ‘idmapping_selected.tab.gz’ sheet available from UniProt to those present in Baktfold’s Swiss-Prot database (n=550,316). We then computed the baseline frequency of GO terms across Swiss-Prot by counting the number of unique proteins linked to that GO term and dividing by 550,316. We then considered every GlobDB CDS that Baktfold could annotate with a Swiss-Prot accession. For all CDSs whose annotated Swiss-Prot accession had at least one linked GO term, we counted the number of unique CDS that GO term was found. We then divided this by the total number of CDS whose annotated Swiss-Prot accession had at least one linked GO term (n=91,887,295 for bacteria and n=14,032,449 for archaea) for bacteria and archaea respectively, yielding the proportion of CDS annotations each GO term was found in. The fold change in GO term presence was calculated by dividing the proportion of CDS annotations each GO term was found in the Baktfold annotations by its baseline frequency in Swiss-Prot. To focus only on GO terms that were reasonably common, we considered only GO terms found in at least 20 proteins in Swiss-Prot.

#### Baktfold Comparison between ProstT5, ESMFold and ColabFold Generated Structures

We used 360 bacterial isolate genomes from NCBI GenBank (‘genbank’ dataset) along with 197 metagenome assembled genomes (‘mag’ dataset) from the Bakta manuscript[3]. We ran Bakta v1.11.4 to re-annotate these genomes given the improvements in Bakta since publication. We then extracted all hypothetical proteins and ran ColabFold v1.5.5[27] implementing AlphaFold2[19] to generate protein structure predictions for all proteins under 3,000 amino acids in length (this being approximately the memory limit of our available GPUs, AMD MI250x on Setonix at the Pawsey Supercomputing Research Centre). Specifically, MSAs were created using both the uniref2302_30 and colabfold_envdb_202 108_db databases using MMseqs2[8] v71dd32, and AlphaFold2 was run in batch mode with one model (‘–num-models 1’) and the default three recycles, without AMBER relaxation. Overall, this yielded 131,027 structures for the genbank dataset and 78,674 structures for the mag dataset. As a protein-language model-based benchmark, additionally, all structures that fit in memory (less than ∼900 amino acids) were then predicted using ESMFold[26], yielding 129,074 structure predictions for the genbank dataset and 76,458 for the mag dataset All Baktfold benchmarking runs were conducted with the ‘Baktfold compare’ command on a HPC node with NVIDIA A100 40GB GPU and 16 threads, utilising Foldseek’s GPU accelerated prefilter[44] via Baktfold’s ‘--foldseek-gpù parameter.

#### Archaea Custom Database and Curation Analysis

We split the Baktfold archaeal genome annotation with custom database annotation benchmarking into two stages. We first built a conservative baseline database of archaeal proteins to see how much, if any, improvements could be observed by introducing a user curated library of annotated proteins into the annotation pipelines. Second was defining a smaller subset of archaeal genomes for benchmarking Baktfold annotation performance for samples across different evolutionary clades.

Our additional annotation protein library is based on GTDB release r226[45], starting with 17,246 archaeal genomes in the release. We successively filtered out single cell and metagenome based entries and genomes with: one or more gaps; one or more ambiguous bases; checkM2[46] completeness scores below 70% and contamination scores above 15%. The remaining archaeal GTDB genomes were then re-queried against most recent NCBI metadata to filter out any assemblies with atypical or suppressed tags, giving us a final archaeal genome set of 1,155 assemblies representing 10 phyla and 226 genera. These 1,155 genomes yielded 3.29 million proteins. After deduplication, the remaining hypothetical/unannotated proteins were removed, leaving 1,993,306 archaeal proteins set to be used as a custom database for annotation.

For the benchmark archaeal genome set our priority was keeping the number of target genomes manageable while ensuring a good degree of genomic and taxonomic variability among the samples. We decided to derive 50 target genomes from our GTDB[45] protein donors to act as control, and added 50 more genomes from GlobDB[41] r226 set to represent diverse, edge case, niche and underrepresented archaeal genomes.

14,839 archaeal genomes from Globdb r226 data were successively filtered, selecting for genomes with 90% or higher CheckM2 completeness score and 10% or less CheckM2 contamination score, then GTDB represented duplicate genomes were removed from the set, leaving us with 1,974 GlobDB based genomes.

Separate phylip distance matrices were built from each representative archaeal genome set from GTDB (1155 genomes) and GlobDB (1,974 genomes) using Skani[47], which were then imported into Julia and processed into 50 UPGMA clusters per data group based on average linkage using Clustering.jl module. A medoid genome was chosen from each of the resulting clusters yielding 100 final representative benchmark archeal genomes, 50 from each of GTDB and GlobDB.

The representative genomes set was processed in Bakta v1.11.4[3], Prokka v1.14.6[4] with default database, Prokka v1.14.6 with our custom protein database, Baktfold v0.1.0 with default database, and Baktfold v0.1.0 with the custom protein database. All representative genomes were then visualised using an unrooted tree build using the GToTree[48] pipeline utilizing a 76 Archaeal single copy gene marker set, with iTOL[49] used to visualize and align plots of annotation percentage next to the tree nodes.

#### Eukaryotic Genome Analysis

Two datasets were used to benchmark Baktfold on micro-eukaryotes. The first was 241 protist genomes from Ensembl protists release 62[50]. Specifically, 802 GenBank format reference genome annotation files (some genomes consisted of multiple GenBank files across different chromosomes) were downloaded from ftp.ebi.ac.uk/ensemblgenomes/pub/release-62/protists, along with taxonomic information (taxonomic identifiers and higher level protist classifications). These were then converted into the required JSON format using the ‘baktfold convert-euk’ command. Baktfold was then run using ‘baktfold run’ with default parameters other the ‘--euk’ flag to indicate a eukaryotic genome. The number of GO terms was calculated by scanning the reference GenBank files. If a CDS had at least one GO term, it was counted.

The second dataset consisted of 713 environmental eukaryotic plankton single-cell (‘SAG’, n=30) and metagenomic assembled genomes (‘MAG’, n=683) from the Tara Oceans eukaryotic genomes SMAG dataset[51]. As the SMAG genomes were not available in GenBank format, we downloaded the bulk protein sequences from https://www.genoscope.cns.fr/tara/. We also downloaded the eggNOG-mapper[12] annotations performed and made available by the authors as part of that work from the same URL, while taxonomic labels for each genome were taken from Supplementary Table 3 of their manuscript. All 10,207,435 proteins were then annotated using ‘baktfold proteins’ using default parameters. A protein was denoted as having a functional eggNOG-mapper annotation if column 22 (denoting the description) in the output was not blank, regardless of its content.

## Results

### Baktfold’s Annotation Workflow

Baktfold accepts either genome annotations via the JSON format output from Bakta, or a file consisting of protein sequences in the ‘.faa’ format. If a user has genome annotations that do not come from Bakta, Baktfold also provides utility scripts to convert prokaryotic GenBank files from Prokka (’baktfold convert-prokkà) or from other annotation pipelines (genbank_to available at https://github.com/linsalrob/genbank_to) to the Bakta JSON format. Baktfold also includes a parser to convert eukaryotic GenBank files (’baktfold convert-euk’).

Baktfold then extracts all hypothetical proteins and uses the ProstT5 protein language model[28] to rapidly predict each query protein’s Foldseek 3Di token encoding. Structure-based homology searches are then conducted with Foldseek[24] against Baktfold’s databases. Following this, all results are exported in GFF3 and INSDC-compliant[52] flat files as well as comprehensive JSON files facilitating automated downstream analysis. Baktfold annotated genomes can easily be submitted to GenBank via NCBI’s table2asn_GFF tool using Baktfold’s GFF3 and Fasta output files, just like Bakta.

**Figure 1:**
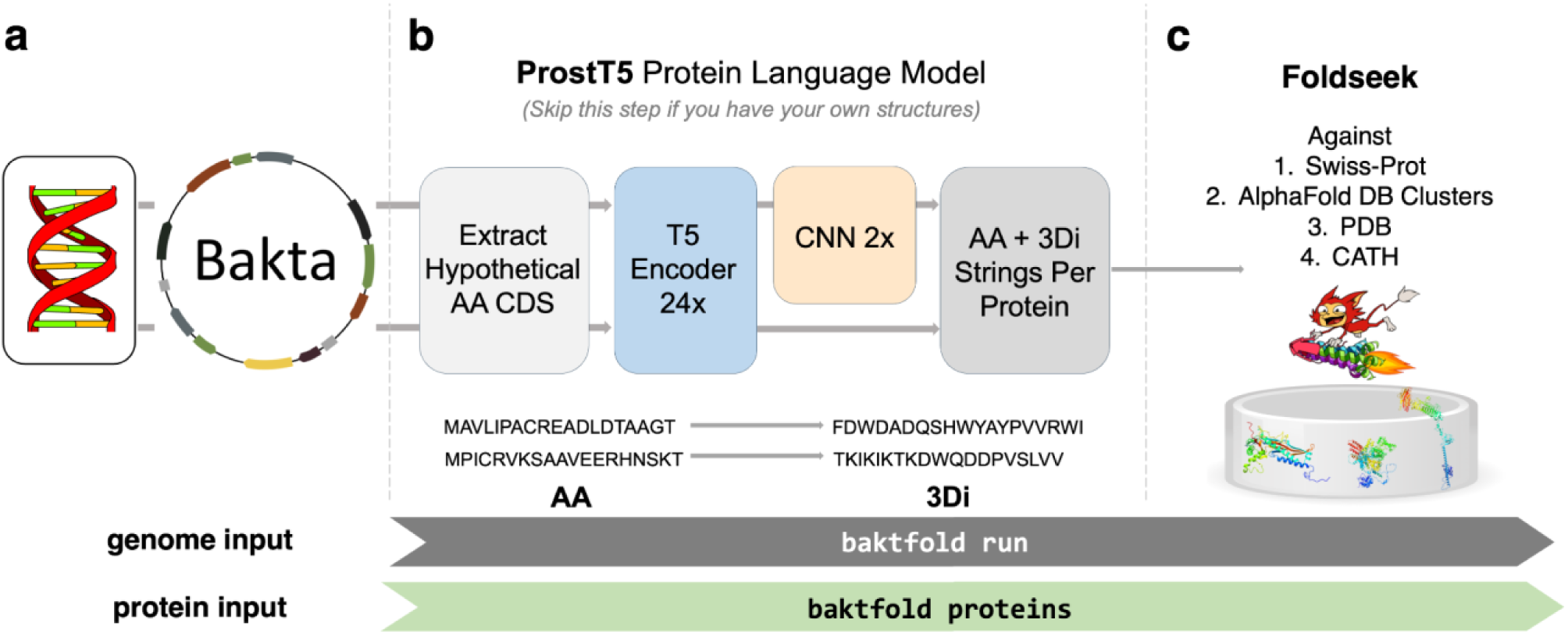
Overview of the Baktfold annotation workflow. (**a**) Baktfold uses either annotated genome input (in Bakta JSON format) or protein multi-FASTA input. For bacteria, we recommend Bakta is run first to predict CDSs and other genomic features such as transfer RNA and transfer-messenger RNAs, then to functionally annotate CDS using sequence-based homology methods. (**b**) Baktfold then predicts Foldseek 3Di tokens for each CDS using ProstT5 pLM’s encoder and CNN. Protein structures can alternatively be used if available. If Bakta genome input is given, Baktfold will only consider unannotated hypothetical proteins to prioritise resource usage on the hardest-to-annotate proteins. (**c**) Baktfold then uses Foldseek to search every CDS using amino acid and ProstT5-predicted 3Di representations against the four databases (Swiss-Prot, AFDB clusters, PDB, CATH) constituting the Baktfold database.

### Baktfold Conducts Comprehensive Sequential Complimentary Database Searches

Taking inspiration from other annotation tools such as Prokka[4] and Pharokka[53], Baktfold uses a series of complimentary databases each with their own strengths to create multiple layers of annotations from different sources.

Baktfold first uses Swiss-Prot (n=590,183)[39], as it provides curated and extremely high-quality annotations. This is followed by a broader search against 3,085,778 non-singleton clusters from AlphaFold Database v6[21, 22]. This is Baktfold’s most computationally demanding step and can be skipped using the ‘--fast’ flag. Final searches are then run against the experimentally resolved Protein Data Bank (PDB)[23] (n=294,848) and domain-level CATH (n=195,223)[40]. The top hit for each database lower than the E-value threshold (default E=1e-03) is then kept for each of the four searches. Baktfold also allows for structure-based searches against custom user-specified Foldseek format databases. If Baktfold finds hits for multiple databases, the overall functional label will be taken preferentially from the user’s custom database (if used), otherwise Swiss-Prot, then AFDB if no Swiss-Prot hits were found, then PDB if neither Swiss-Prot nor AFDB hits were found and finally CATH if no other databases were hit. All top hit accessions are available in ‘baktfold.inference.tsv’ while the detailed foldseek statistics are available for each database in the respective ‘_tophit.tsv’ files in Baktfold’s output.

### Baktfold vastly improves genome annotation across the prokaryotic tree of life

To benchmark the performance of Baktfold across the prokaryotic tree of life, we ran Baktfold across 305,825 species representatives from GlobDB[41], an extensive database of archaeal and bacterial species representatives, along with 8.8M non-redundant plasmid-encoded genes from IMG/PR[42].

Overall, Baktfold annotated a median 87.8% of all CDS across GlobDB bacterial genomes (n=290,259), compared to a median 72.9% with Bakta[3] and 48.1% with Prokka[4] (Supplementary Figure 1), with the median annotation rate of hypothetical proteins (i.e. unable to be annotated by Bakta) of 50.1%. Baktfold annotated fewer hypothetical proteins in genomes that were already well annotated (i.e. >90% annotation rate) by Bakta (Figure 2A-B), though Baktfold was still able to annotate a median of 23.4% of hypothetical proteins even on this extremely challenging set of remaining hypothetical proteins. This was not present when considering Prokka instead of Bakta annotations (Supplementary Figure 1), illustrating the benefit of using Bakta over Prokka for sequence-homology based annotation. On archaea, Baktfold annotates a median 71.5% of CDS (n=14,812) compared to 35.8% with Prokka and 10.3% with Bakta (n.b. Bakta was not designed for archaeal genome annotation). 68.0% of hypothetical proteins were able to be annotated by Baktfold on the median archaeal genome.

**Figure 2:**
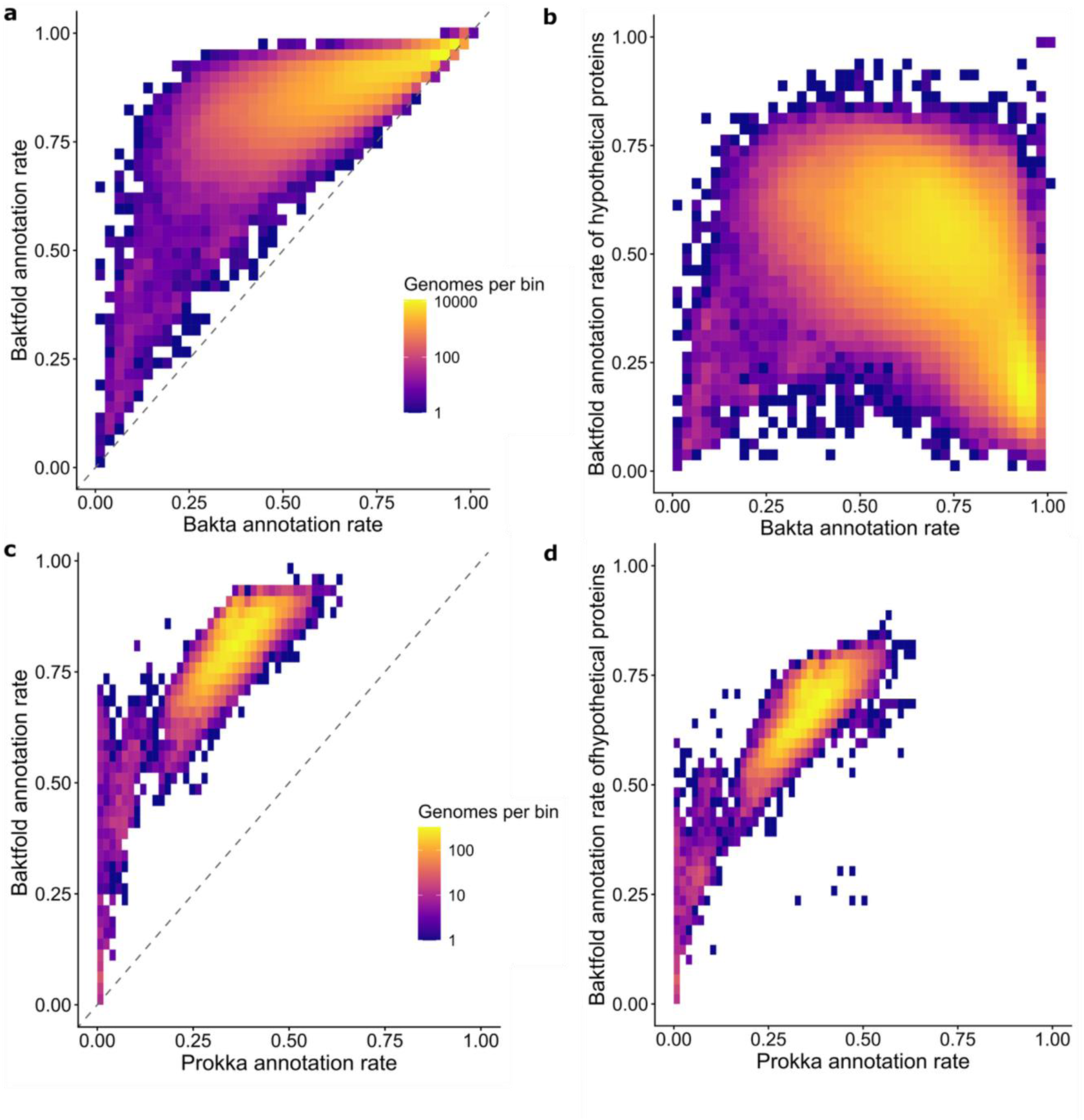
Baktfold annotates hypothetical proteins across the bacterial and archaeal kingdoms. (a) Binned count heatmaps showing the number of bacterial genomes in GlobDB (n=290,258) of Baktfold overall annotation rate (y-axis) compared to Bakta (x-axis) (b) the same bacterial genomes with Baktfold’s annotation rate of hypothetical proteins only on the y-axis (c) binned count heatmaps showing the number of archaeal genomes in GlobDB (n=14,058) for each bin of Baktfold overall annotation rate (y-axis) compared to Prokka (x-axis) and (d) the same archaeal genomes with Baktfold’s annotation rate of hypothetical proteins only on the y-axis.

Baktfold annotated 79.0% of all non-redundant plasmid proteins from IMG/PR, compared to 70.2% for Bakta, yielding an annotation rate of 29.5% of those difficult proteins unable to be annotated by Bakta. Baktfold showed improvement for longer plasmid proteins (100 amino acids and longer in length), with Baktfold able to annotate 46.5% of the approximately 1.5M proteins that Bakta could not, while for short proteins (<100AA), Baktfold added far fewer additional annotations, annotating only 6.5% of the approximately 1.1M short proteins (Supplementary Figure 2).

While benchmarking functional annotation is extremely challenging, especially where the added annotations are further into the twilight zone of sequence similarity[25] often referred to as “microbial dark matter”[13], we sought to understand whether Baktfold’s annotations seem generally reasonable. To do this, we looked at the proportion of GO terms linked to CDS annotated by Baktfold with Swiss-Prot accession across GlobDB and compared them to their baseline frequency in the Swiss-Prot database. Two of the top five GO terms that are disproportionately present in bacteria (Figure 2E) relate to Type IV pilus structure and motility[54] (Figure 3A), while two relate to specific transferases and the other transducer activity. For archaea, the top two hits related to archaea flagellum[55] and S-layer organisation[56], while the others related to synthase and reductases (Figure 3B).

**Figure 3:**
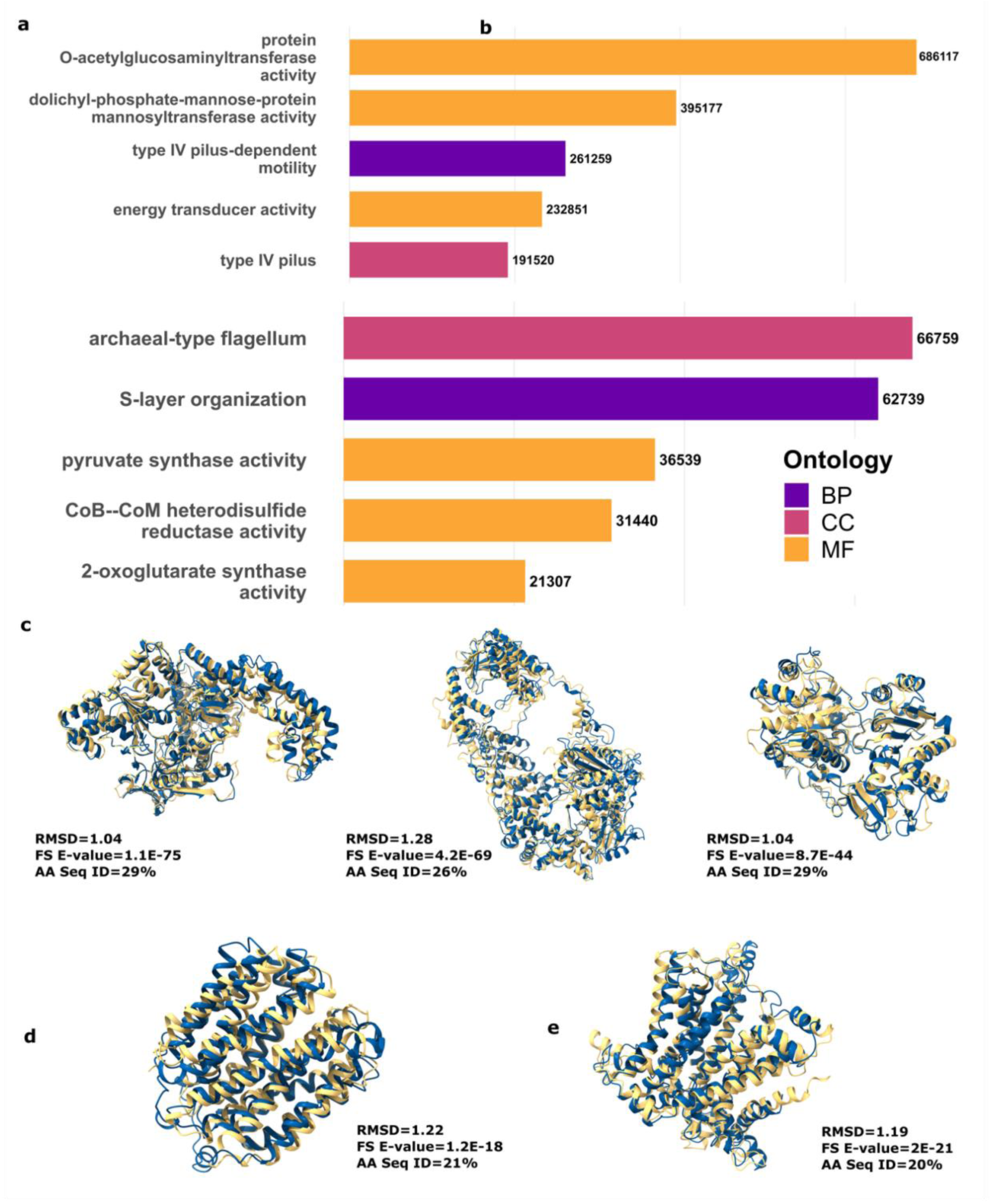
Baktfold enables detailed functional annotation into the twilight zone of sequence similarity. (a-b) The top 5 GO terms that are disproportionately present in Baktfold CDS Swiss-Prot annotations compared to the baseline frequency across Swiss-Prot across bacteria (1) and archaea (b) genome from GlobDB. The number of CDS each GO term occurs in across all GlobDB genomes is marked on the right of each bar. BP = biological process, CC = cellular component and MF = molecular function. (c) Structural superposition of each subunit of the putative RecBCD enzyme locus compared to Swiss-Prot best hit found in Escherichia coli. Yellow shows the ColabFold structure of the CDS from GCA_036689255, while blue shows the Swiss-Prot entry. (d) Structural superposition of the annotated inner membrane metabolite transport protein in GCA_023256405 with the Swiss-Prot best hit Q13I96 found in Paraburkholderia xenovorans. (e) Structural superposition of the annotated Ferric transport system permease protein FbpB in GCA_023256405 with the Swiss-Prot accession Q44123 found in Actinobacillus pleuropneumoniae.

To illustrate Baktfold’s ability to annotate hypothetical CDS deep into the twilight zone of sequence identity (i.e. between 20-35% sequence identity), we took two GlobDB genomes where Baktfold had amongst the highest rate of hypothetical protein annotations and inspected the annotations of selected CDS (bacteria: NCBI accession GCA_036689255 *Candidatus stammera capleta*, archaea: NCBI accession GCA_023256405 *Nitrososphaerales* archaeon). For example, on bacterium GCA_036689255, Baktfold confidently annotated a putative operon homologous to all three units of the RecBCD enzyme Exodeoxyribonuclease V. All three monomer subunits have low sequence identity (<30%) but extremely strong structural similarity to the respective RecBCD subunit (Figure 3C). On the archaeon GCA_023256405, Baktfold was able to annotate an inner membrane metabolite transport protein Ygcs with 21% sequence identity and E-value of 1.2e-18 and RMSD of 1.22 to AFDB/Swiss-Prot accession Q13I96 (Figure 3D) and a Ferric transport system permease protein FbpBn with 20% sequence identity and E-value of 2e-21 and RMSD of 1.19 to AFDB/Swiss-Prot accession Q44123 (Figure 3E).

### Baktfold with ProstT5 enables annotation in minutes and is almost as sensitive as using AlphaFold2

It has been previously shown that using ProstT5 for Foldseek 3Di token inference instead of leading protein structure prediction tools ESMFold[24] or ColabFold/AlphaFold2[19, 27] structures is orders of magnitude more computationally efficient[28, 57]. It has also been shown this nonetheless yields comparable performance across remote structural homology benchmarks[28] and bacteriophage protein annotation tasks[57]. Following this, we then compared Baktfold’s annotation performance using the ProstT5 protein language model against using ColabFold/AlphaFold2 and ESMFold. We used 360 bacterial isolates from NCBI Genbank (‘Genbank dataset’) along with 197 metagenome assembled genomes (‘mag dataset’) from Bakta[3] and ran Bakta followed by Baktfold with three different structure information prediction tools: ProstT5 (Baktfold’s default), ESMFold and ColabFold/AlphaFold2 and otherwise identical parameters.

The runtime of Baktfold with ProstT5 was linearly correlated (r=0.97) with the number of input hypothetical proteins from Bakta to be annotated (Figure 4A). With GPU acceleration, Baktfold completed in between 30 seconds (n=101 hypothetical proteins) and 457 seconds (n=1989 hypothetical proteins) of wallclock runtime across the MAG dataset (Figure 4A). On the benchmarked hardware (1x NVIDIA A100 40GB GPU, 16 CPU cores), the Bakta and Bakftold combined wallclock runtime ranged between 383 seconds and 1218 seconds across the mag dataset (Supplementary Table 1).

**Figure 4:**
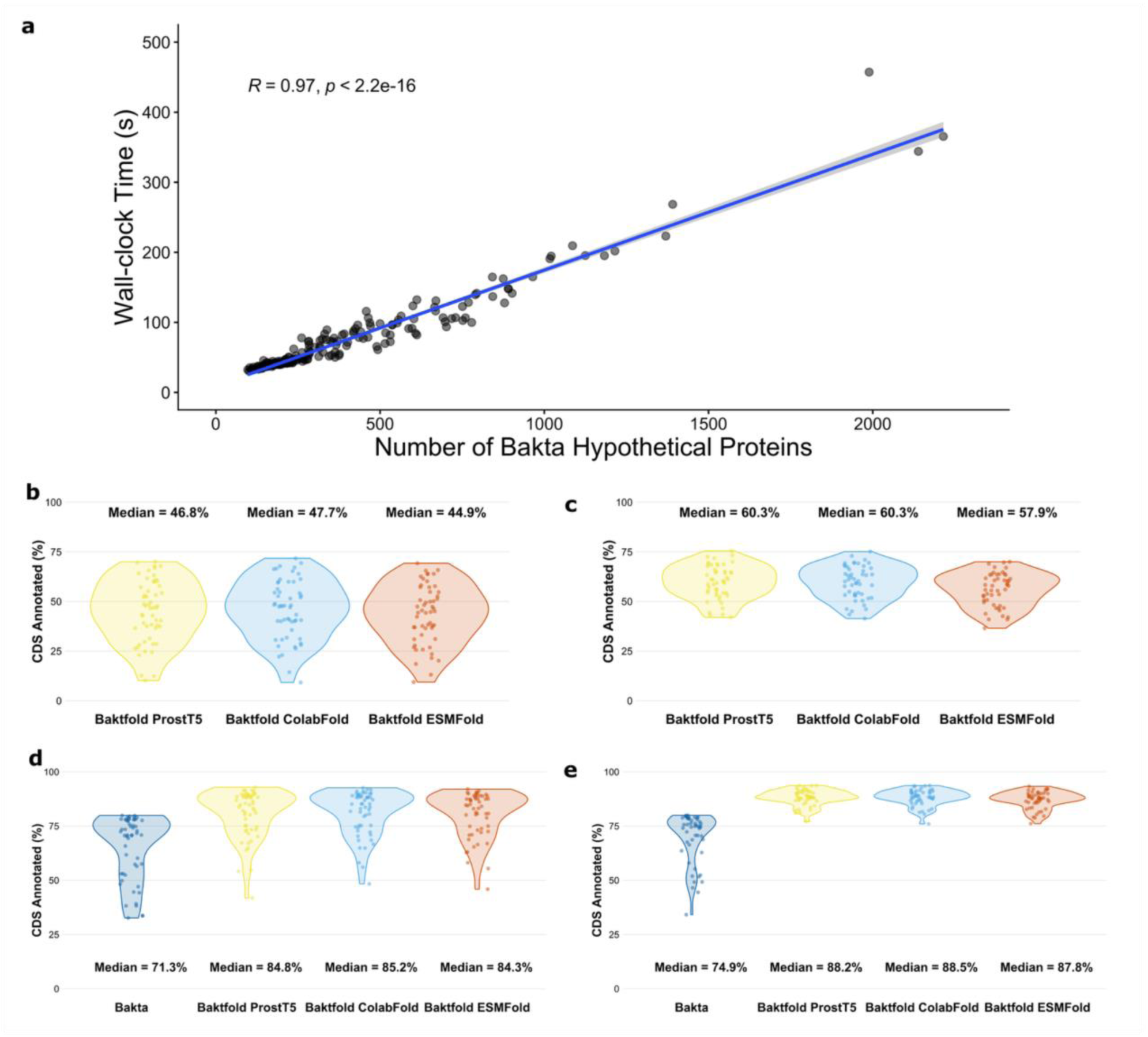
Baktfold’s performance with ProstT5 is similar to full structure prediction models though taking only minutes to complete. (a) Baktfold’s wallclock runtime (y-axis) compared to the number of input hypothetical proteins from Bakta to be annotated (x-axis) for 197 genomes from the magdataset. (b-c) Baktfold’s hypothetical CDS annotation rate using ProstT5, ColabFold and ESMFold for structural information inference for Genbank (n=55, b) and mag (b=49, c) dataset genomes where Bakta functionally annotated less than 80% of CDS. (d-e) The overall annotation of the same genomes on the Genbank (d) and mag (e) datasets.

Considering only the hardest genomes in both datasets (i.e. where Bakta could annotate fewer than 80% of CDS), Baktfold with ProstT5 was almost identical in terms of annotation rate of hypothetical proteins (median 46.8% Genbank; 60.3% mag) compared to Baktfold with ESMFold (median 44.9% Genbank; 57.9% mag) and ColabFold/AlphaFold2 (median 47.7% Genbank; 60.3% mag) across both Genbank (Figure 4B) and mag (Figure 4C) datasets. Overall, Baktfold with any of the structural prediction tools yielded median annotation rates of over 90% on both datasets on these difficult genomes (Figure 4D-E). Similar performance was observed across all benchmarked genomes, including those where Bakta could annotate over 80% of CDS (Supplementary Figure 3).

### Structure-informed annotation with Baktfold revolutionises archaeal annotation

Given Baktfold’s strong performance across archaea and to showcase the custom database functionality of Baktfold, we decided to conduct a deeper archaea-specific analysis. Using a curated phylogenetically diverse archaeal genome benchmarking set of 100 archaeal genomes, Baktfold outperformed all other annotation pipelines using default databases across all surveyed taxonomic groups (Figure 5A), showing a median annotation rate of 72.71% compared to Prokka’s 38.85% (and Bakta’s 10.04%). Using our additional customised archaeal protein database comprising over 1.99 million archaeal proteins (see methods) increased the median annotation rate to 85.61%, with consistent improvements across taxa (Figure 5B).

**Figure 5:**
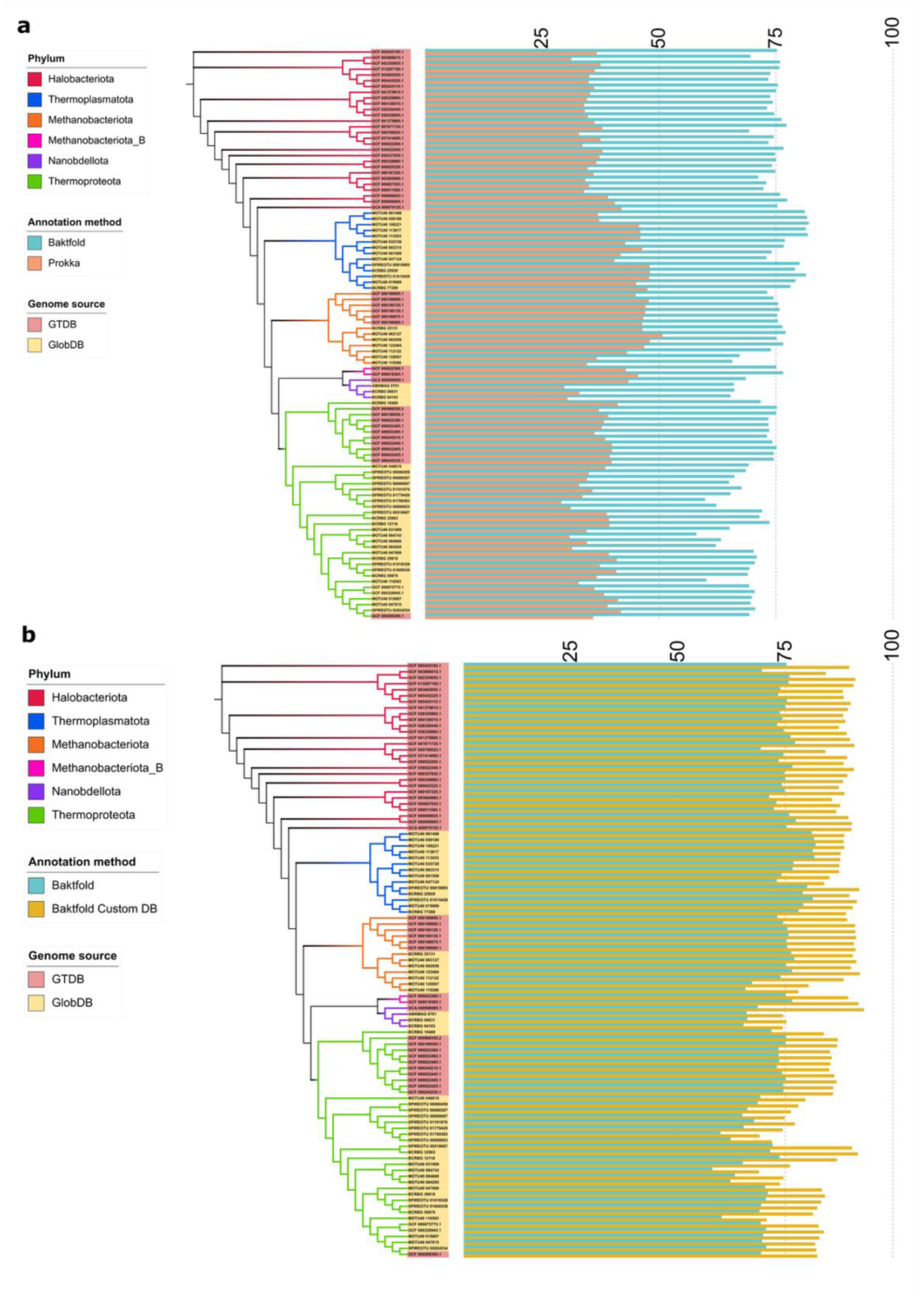
Baktfold excels on archaeal annotation across taxa and is augmented by archaeal-specific custom databases. (a-b) Cladograms for 100 curated archaeal genomes from GTDB and GlobDB (left) with the functional annotation rate presented as bars to the right of each genome for (a) Baktfold and Prokka (both) default databases and (b) Baktfold and Baktfold with our customised archaeal protein database consisting of nearly 2M archaeal proteins.

While the customised database closes the annotation gap between Baktfold and Prokka, Baktfold still maintained a noticeable lead of 85.61% average annotation versus 79% of Prokka (Supplementary Figure 4), suggesting that structure-based homology approaches remain more sensitive even where targeted, customised databases of interest are used to augment genome annotation. The annotation rate for genomes from some taxonomic groups such as *Thermoplasmatota* and *Thermoproteota* remains low (default 36.95% and with custom database 49.23%) using Prokka’s sequence-homology based annotation methods (Figure 5B). Baktfold, however, maintains a significantly higher annotation rate of 81.16% using its default database and 88.72% using our custom database, suggesting that for certain archaeal taxa, structure-based homology methods far outperform sequence alignment-based annotation.

### Baktfold can rapidly and consistently annotate micro-eukaryotes using structural homology

Though designed with bacteria and archaea in mind, given the taxonomic breadth of the constituent databases, Baktfold can also be applied for fast functional annotation of micro-eukaryotes. To illustrate this functionality, we tested Baktfold’s performance across two datasets: 241 genomes comprising of 196 distinct taxonomy IDs with manually curated assemblies and genome annotations from Ensembl Protists[50] (release 62) and over 10.2 million CDS taken from 713 environmental eukaryotic plankton single-cell (‘SAG’, n=30) and metagenomic assembled genomes (‘MAG’, n=683) from the SMAG dataset[51] that had been functionally annotated using sequence-based homology methods (combined with orthology assignments) with eggNOG-mapper[12] as part of that study.

Across all EnsemblProtists genomes, Baktfold was able to functionally annotate 70.0% of all CDS (n=1,860,459/2,657,255), larger than the 60.7% (n=1,613,095/2,657,255) of CDS with Gene Ontology (GO) term annotations in the reference genome annotations. As these represent the set of currently best annotated protist genomes, this suggests Baktfold provides substantial improvement in inferring gene function in protists.

The improvement was not consistent across all protist taxa (Figure 6). Across the higher classification levels provided by Ensembl (Figure 6A), Baktfold performed strongest on *Amoebozoa* (77.1% total functional annotation rate) and *Stramenopiles* (73.5%), with the biggest difference between Baktfold and GO annotation rates occurring for *Stramenopiles* (12.9%). Baktfold performed less well on *Rhizaria* (61.0%) and *Fornicata* (63.9%), while *Alveolata* showed the smallest difference between Baktfold and GO (3.5%). At the NCBI taxonomic identifier (‘taxon ID’) level, Baktfold provided more functional annotations than existing GO terms for 78.1% (153/196) of taxa (Figure 6B). Slime mold *Tieghemostelium lacteum* (taxon ID 361077) had the highest Baktfold annotation rate (86.6%), while the plant pathogen *Globisporangium ultimum* (taxon ID 1223559) had the highest difference between Baktfold and GO terms (37.3% GO, 69.3% Baktfold). *Alveolata* was the notable group where Baktfold was relatively less effective, with 40.4% (38/94) taxon IDs having fewer functionally annotated CDS with Baktfold than GO terms, while each taxon in *Stramenopiles* (n=44), *Amoebozoa* (n=15) and *Fornicata* (n=6) had a higher annotation rate with Baktfold than GO terms.

**Figure 6:**
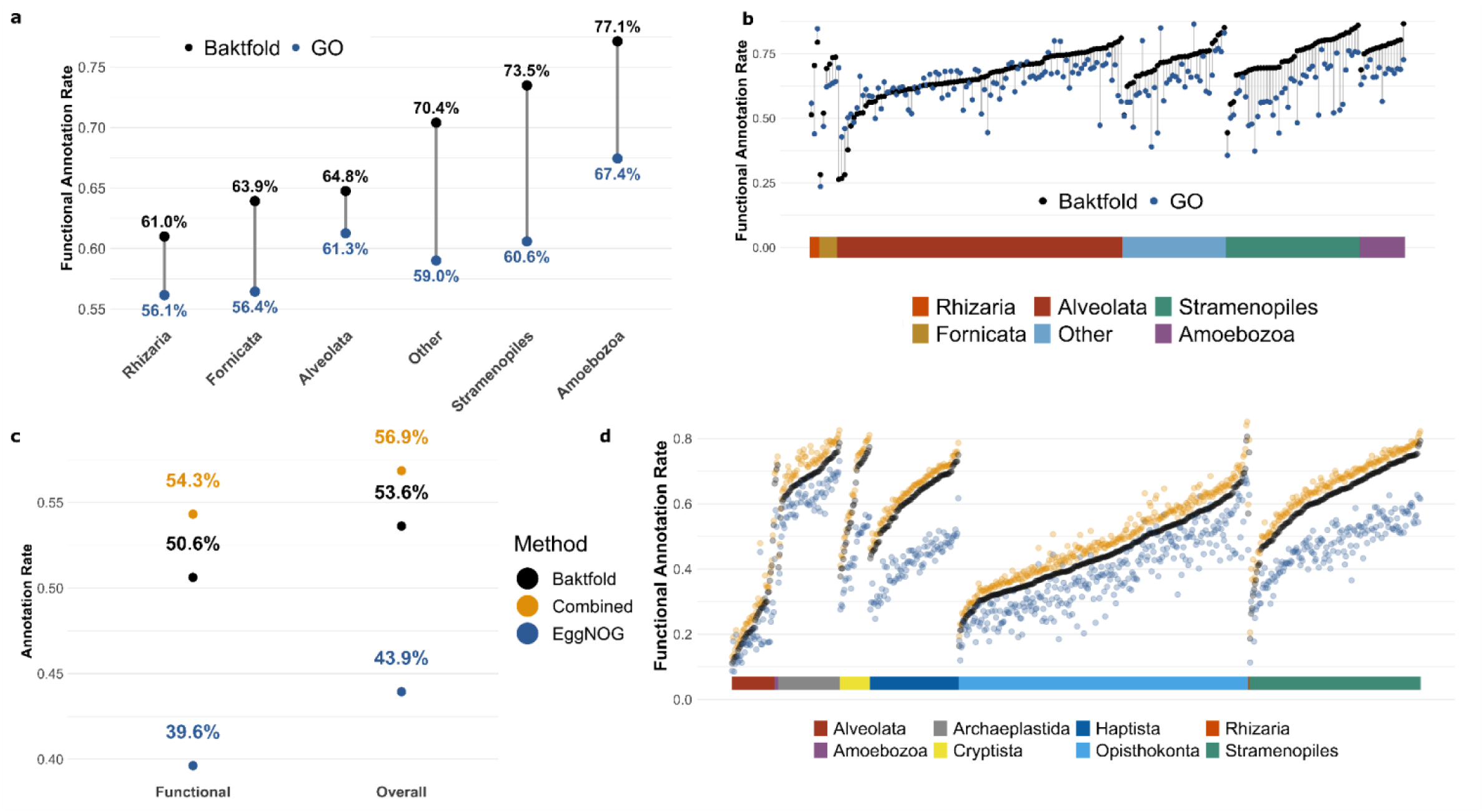
Baktfold annotation performance across selected protist genomes. (a) Functional CDS annotation rate (black) compared to the proportion of Ensembl Protists reference genome CDS with at least one Gene Ontology (GO) term annotation (blue) across 241 genomes, calculated within each protist classification. (b) Same as (a) but calculated across each protist taxon ID in the dataset. (c) Functional and overall annotation rates of Baktfold (black), eggNOG-mapper (blue) and combined (orange) across 713 MAG and SAG eukaryotic plankton genomes from the SMAG dataset (d) Same as (c) functional annotation rates for each method calculated across each genome in the dataset, sorted in ascending Baktfold annotation rate per taxonomic super group.

For the SMAGs, Baktfold had an overall CDS functional annotation rate of 50.6% (n=5,167,383/10,207,435) and an overall annotation rate (i.e. including hits to database proteins with no functional annotation) of 53.6% (5,473,475/10,207,435). By contrast, the respective annotation rates were 39.6% and 43.9% for eggNOG-mapper (Figure 6C), while the combined annotation rates (i.e. combining Baktfold and eggNOG-mapper) were 54.3% and 56.9%, highlighting the complimentary of sequence and structure-based homology detection methods. At the per genome level, 683/713 genomes had higher functional annotation rates with Baktfold than eggNOG-mapper (median 54.8% for Baktfold and 43.3% for eggNOG-mapper), with substantial variation within each taxonomic super group (Figure 6D). Baktfold performed particularly strongly on functional annotation for *Cryptista* and *Hapista* genomes compared to eggNOG-mapper (overall, 61.8% vs 46.7% for *Cryptista* and 63.9% vs 46.7% for *Hapista*, respectively) (Supplementary Figure 5A). MAGs generally higher functional annotation rates than SAGs regardless of method (Supplementary Figure 5B) (Baktfold functional CDS annotation rate of 51.1% for MAG vs 42.1% for SAG, compared to 40.1% and 30.3% for eggNOG-mapper), suggesting the impacts of lower quality SAG assemblies did not disproportionately impact Baktfold compared to eggNOG-mapper, or vice versa.

## Discussion

While large protein structure databases have been widely available and searchable for a few years, there exists no automated and easy-to-use tool that makes use of protein structural information to annotate microbial genomes. While well-studied microbes such as *Escherichia coli* and *Staphylococcus aureus* are well annotated using sequence-based homology approaches[1], much microbial dark matter remains to be functionally illuminated[13].

In this study, we present Baktfold, our tool that uses pLM (ProstT5) based structural information in conjunction with Foldseek to perform sequential searches across well-annotated protein structure databases. We show that Baktfold improves functional annotations across the tree of microbial life, including bacteria, plasmids, archaea and micro-eukaryotes. More detailed analyses of selected Baktfold annotations reveal that Baktfold is able to detect strong structural homologs deep into the twilight zone of sequence alignment[25]. As such, Baktfold is an essential tool to enrich protein annotations in conjunction with existing sequence-based homology genome annotation across the microbial tree of life. Given Baktfold’s added value in annotating hypothetical proteins in bacteria and plasmids, and particularly in archaea[15], we anticipate it will be of especial interest to researchers from this community. Baktfold will provide the most value in annotating proteins that are currently the hardest to annotate, particularly those longer than 100AA, below which protein structure prediction can become unreliable[58].

Overall, we hope the functional annotations of the hardest microbial proteins provided by Baktfold allows for large-scale hypothesis generation guiding future *in vitro* and *in silico* research into protein and cellular function.

## Data Availability

The Baktfold database is hosted at Zenodo (https://zenodo.org/records/17347516) mirrored on HuggingFace (https://huggingface.co/datasets/gbouras13/baktfold-db). All code and data required to reproduce the analyses in this manuscript are found at https://github.com/gbouras13/baktfold-analysis (code and small data) and https://zenodo.org/records/19333697 (large data).

## Code Availability

Baktfold is open-source software available at https://github.com/gbouras13/baktfold. All other code required to recreate the results in this manuscript can be found at https://github.com/gbouras13/baktfold-analysis.

## Authors and contributors

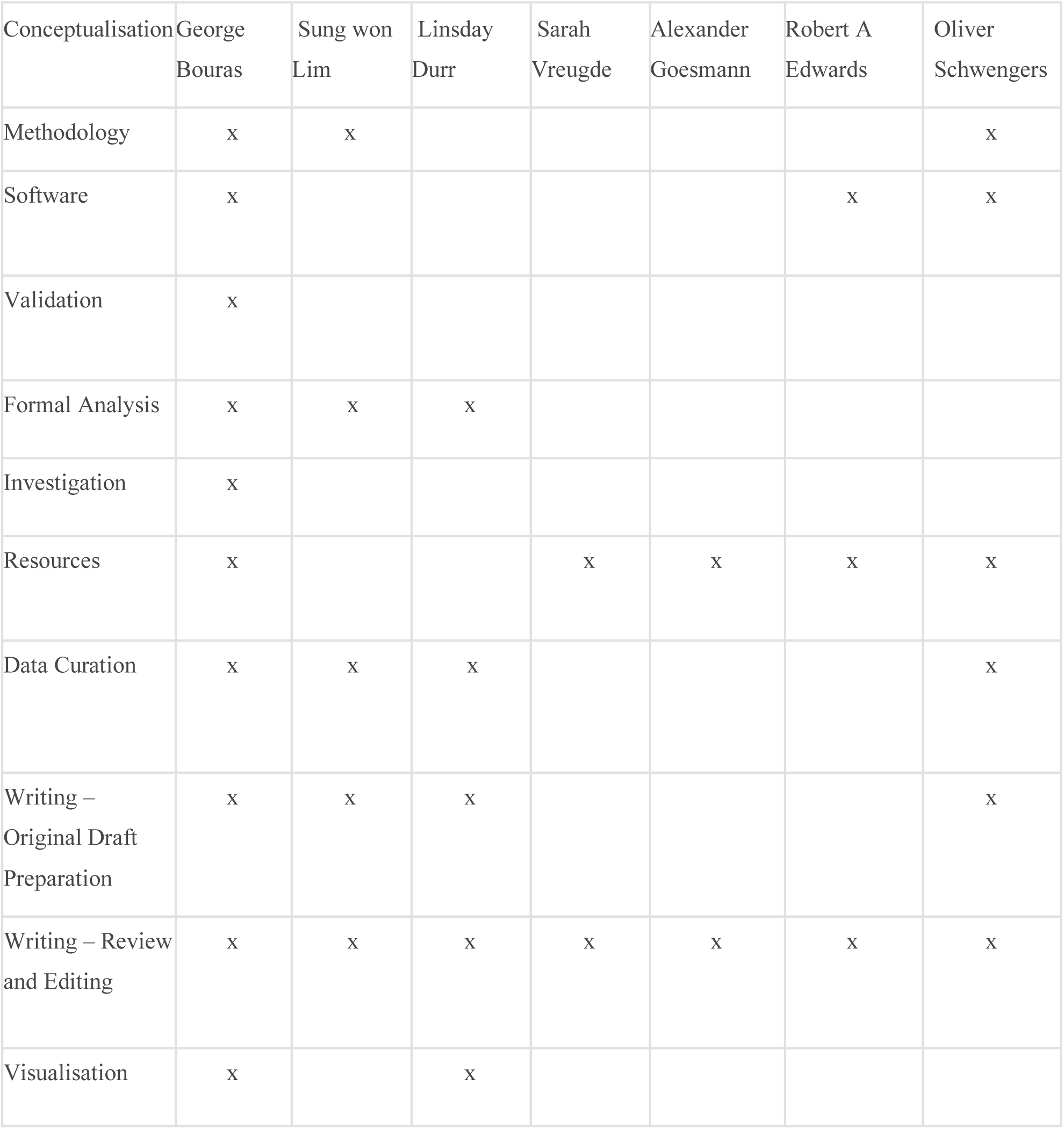
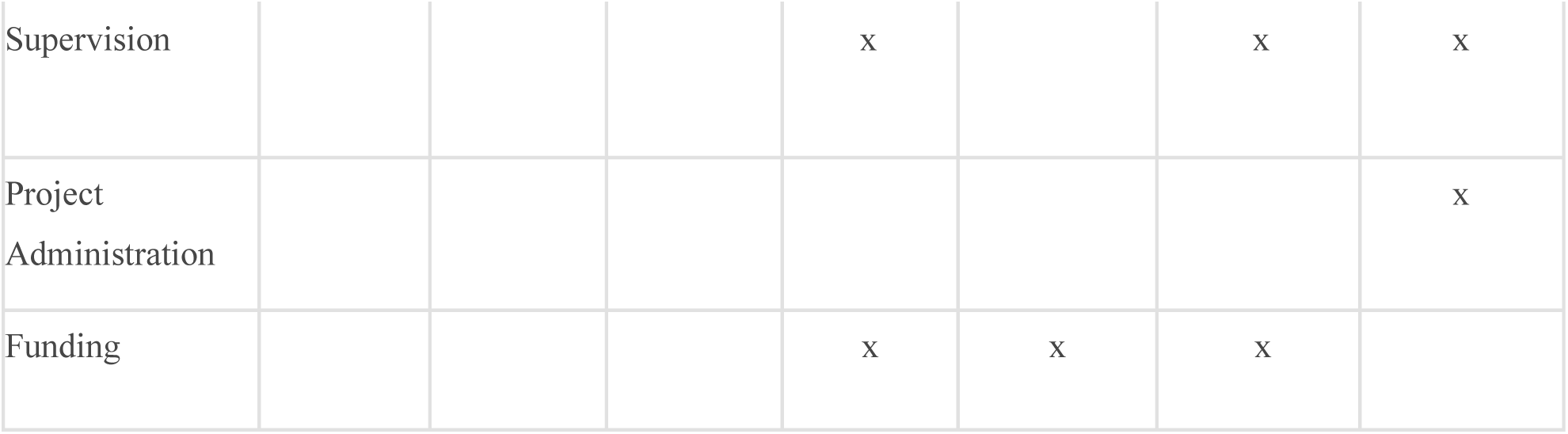

| Conceptualisation | George Bouras | Sung won Lim | Lindsay Durr | Sarah Vreugde | Alexander Goesmann | Robert A Edwards | Oliver Schwengers |
| --- | --- | --- | --- | --- | --- | --- | --- |
| Methodology | x | x |  |  |  |  | x |
| Software | x |  |  |  |  | x | x |
| Validation | x |  |  |  |  |  |  |
| Formal Analysis | x | x | x |  |  |  |  |
| Investigation | x |  |  |  |  |  |  |
| Resources | x |  |  | x | x | x | x |
| Data Curation | x | x | x |  |  |  | x |
| Writing – Original Draft Preparation | x | x | x |  |  |  | x |
| Writing – Review and Editing | x | x | x | x | x | x | x |
| Visualisation | x |  | x |  |  |  |  |
| Supervision |  |  |  | x |  | x | x |
| Project Administration |  |  |  |  |  |  | x |
| Funding |  |  |  | x | x | x |  |

## Acknowledgements

This work was supported with the assistance of resources and services from Phoenix HPC at Adelaide University and Pawsey Supercomputing Research Centre, which is supported by the Australian Government. We would like to thank Fabien Voisin and Sarah Beecroft for their assistance in operating ColabFold at scale at Phoenix and Pawsey, respectively, with extra acknowledgement to Sarah for containerizing ColabFold for use on Setonix’s AMD GPUs. We would also like to thank Renee Green for her assistance in developing the template for Figure 1, Susie Grigson, who provided the idea to specifically analyse plasmids, and Martin Steinegger, Milot Mirdita, Michael Heinzinger and Victor Mihaila for their continued development and maintenance of ProstT5-Foldseek and related discussions and advice.

## Supplementary Figures

**Supplementary Figure 1:**
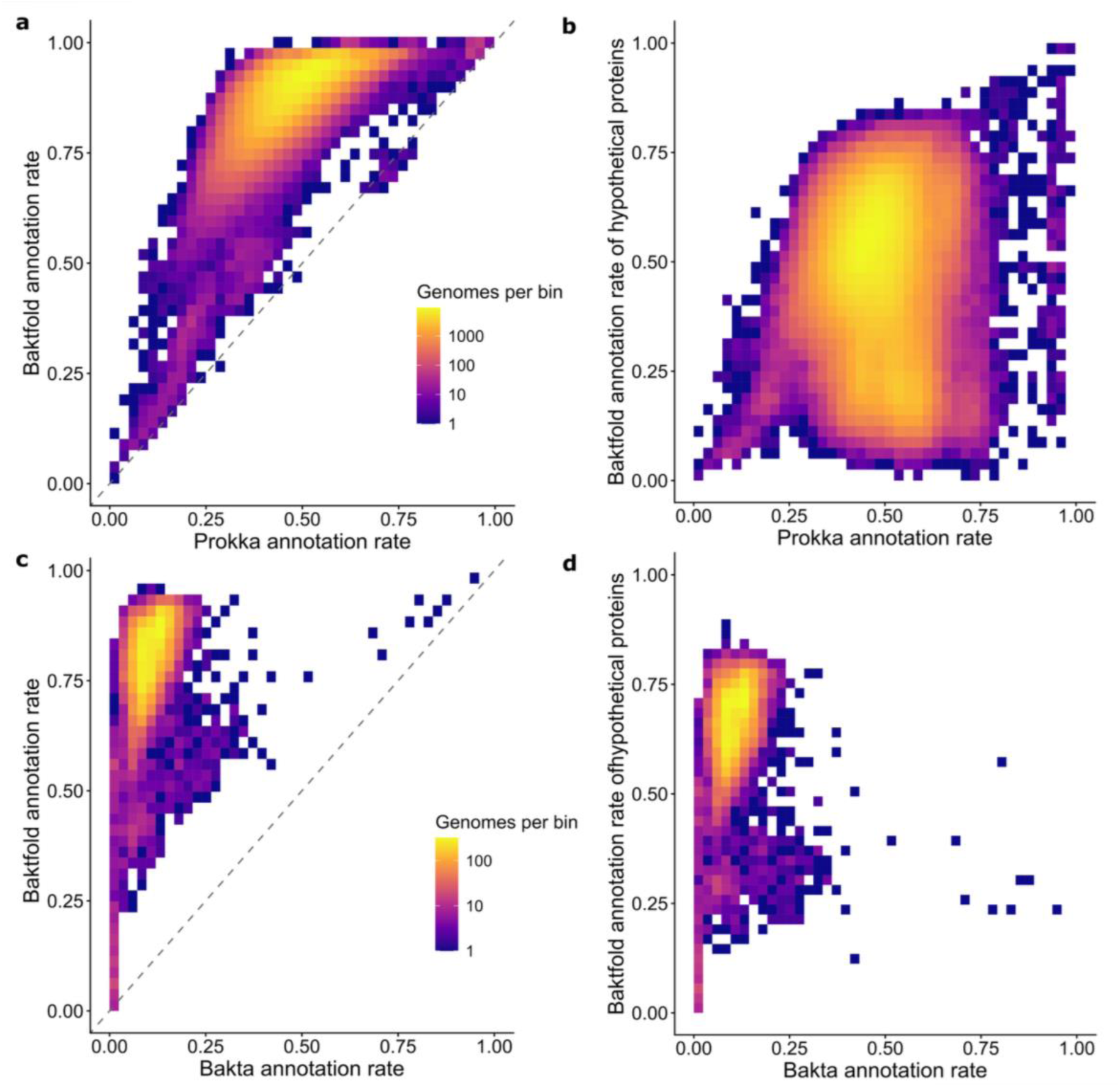
(a) Binned count heatmaps showing the number of bacterial genomes in GlobDB (n=290,258) for each bin of Baktfold overall annotation rate (y-axis) compared to Prokka (x-axis) (b) the same bacterial genomes with Baktfold’s annotation rate of hypothetical proteins only on the y-axis (c) binned count heatmaps showing the number of archaeal genomes in GlobDB (n=14,058) for each bin of Baktfold overall annotation rate (y-axis) compared to Bakta (noting that Bakta was not designed to annotate archaea) (x-axis) and (d) the same archaeal genomes with Baktfold’s annotation rate of hypothetical proteins only on the y-axis.

**Supplementary Figure 2.**
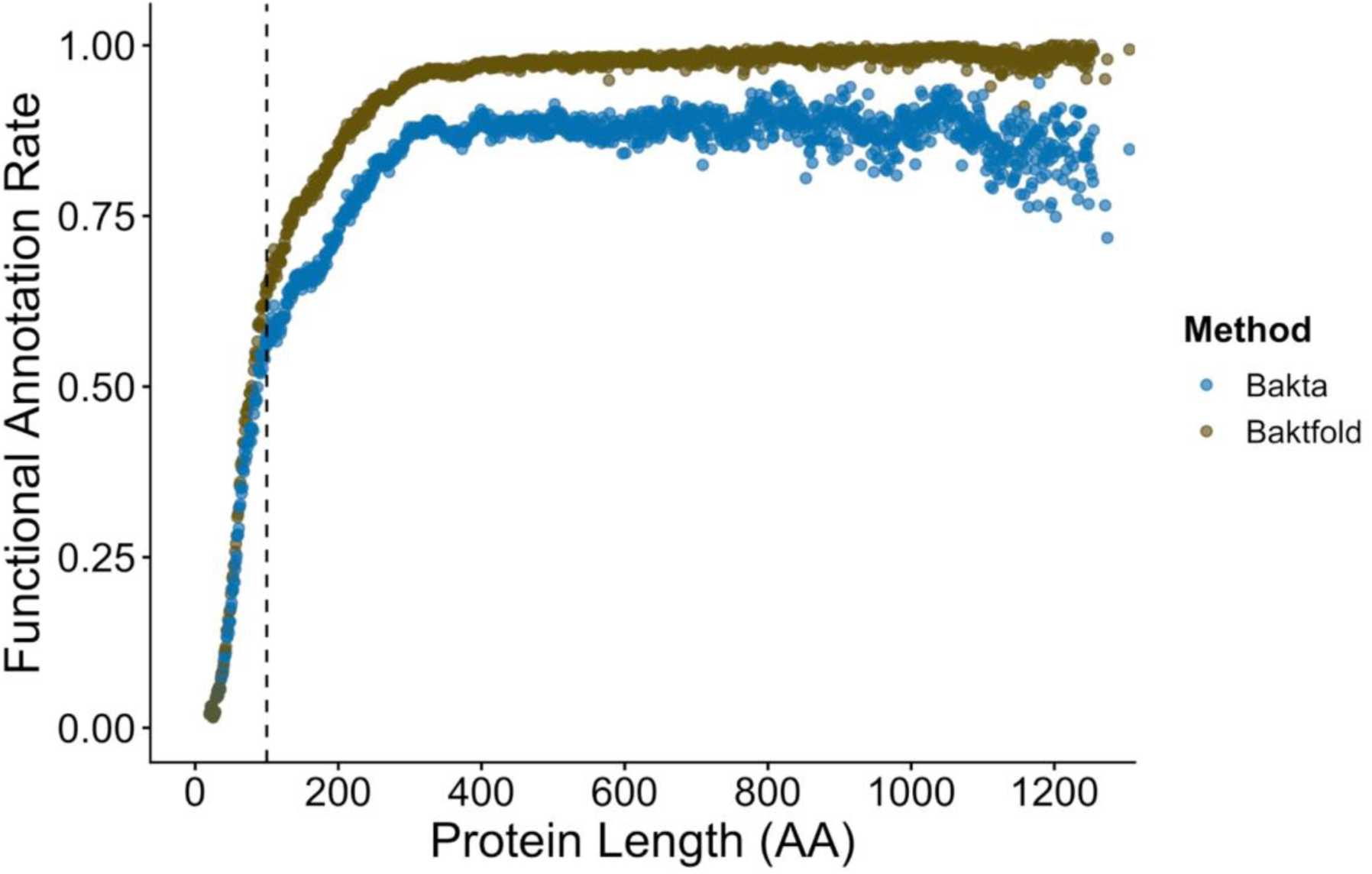
: Functional protein annotation rate for Baktfold (gold) and Bakta (blue) at each length between 20-1,250 amino acids proteins for approximately 8.8M non-redundant plasmid-encoded genes from IMG/PR. The dashed vertical line indicates 100 AA.

**Supplementary Figure 3:**
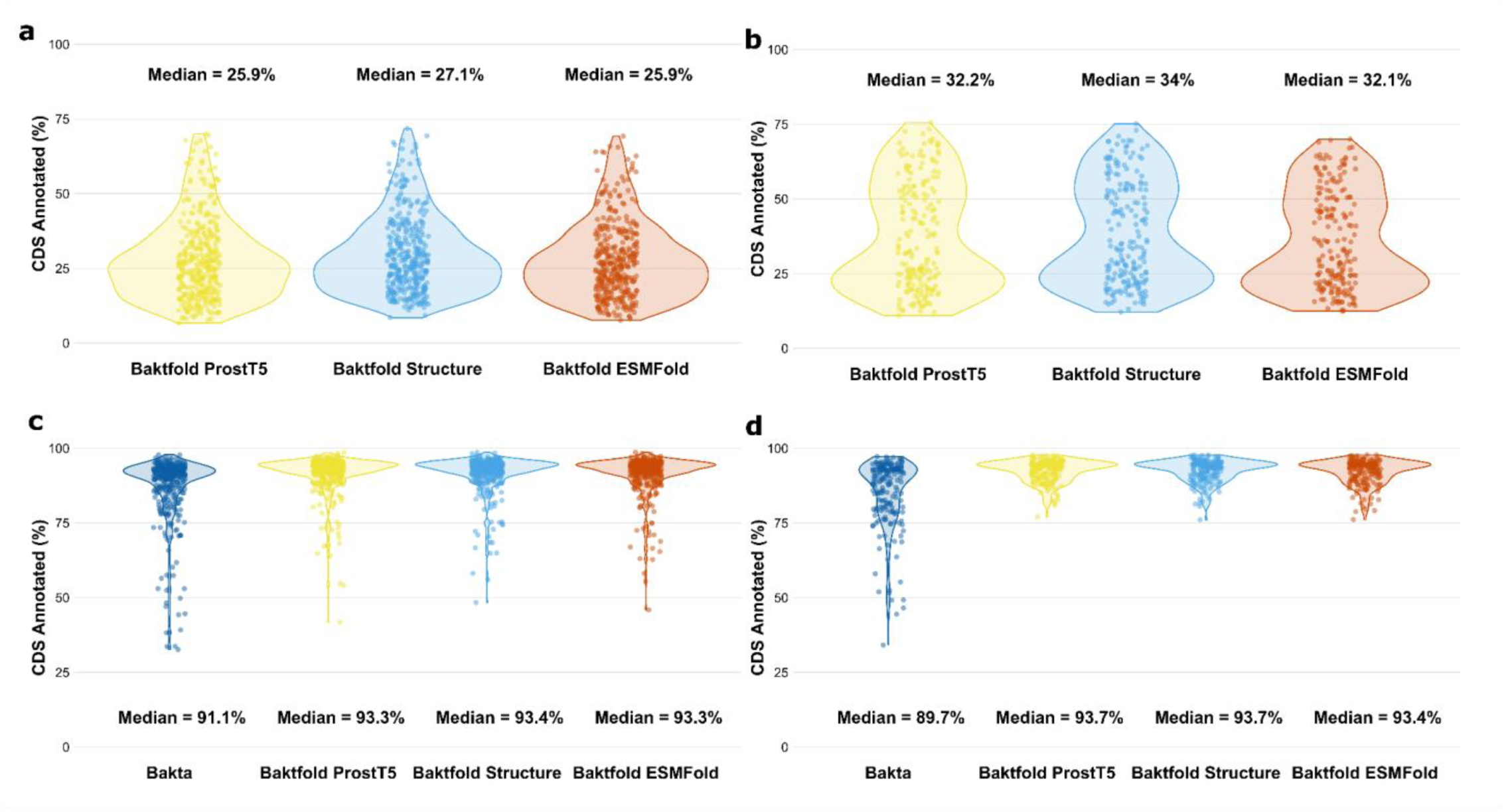
(a-b) Baktfold’s hypothetical CDS annotation rate using ProstT5, ColabFold and ESMFold for structural information inference for Genbank (n=360, b) and MAG (b=197, c) genomes. (c-d) The overall annotation of the same genomes on the Genbank (c) and MAG (d) datasets.

**Supplementary Figure 4:**
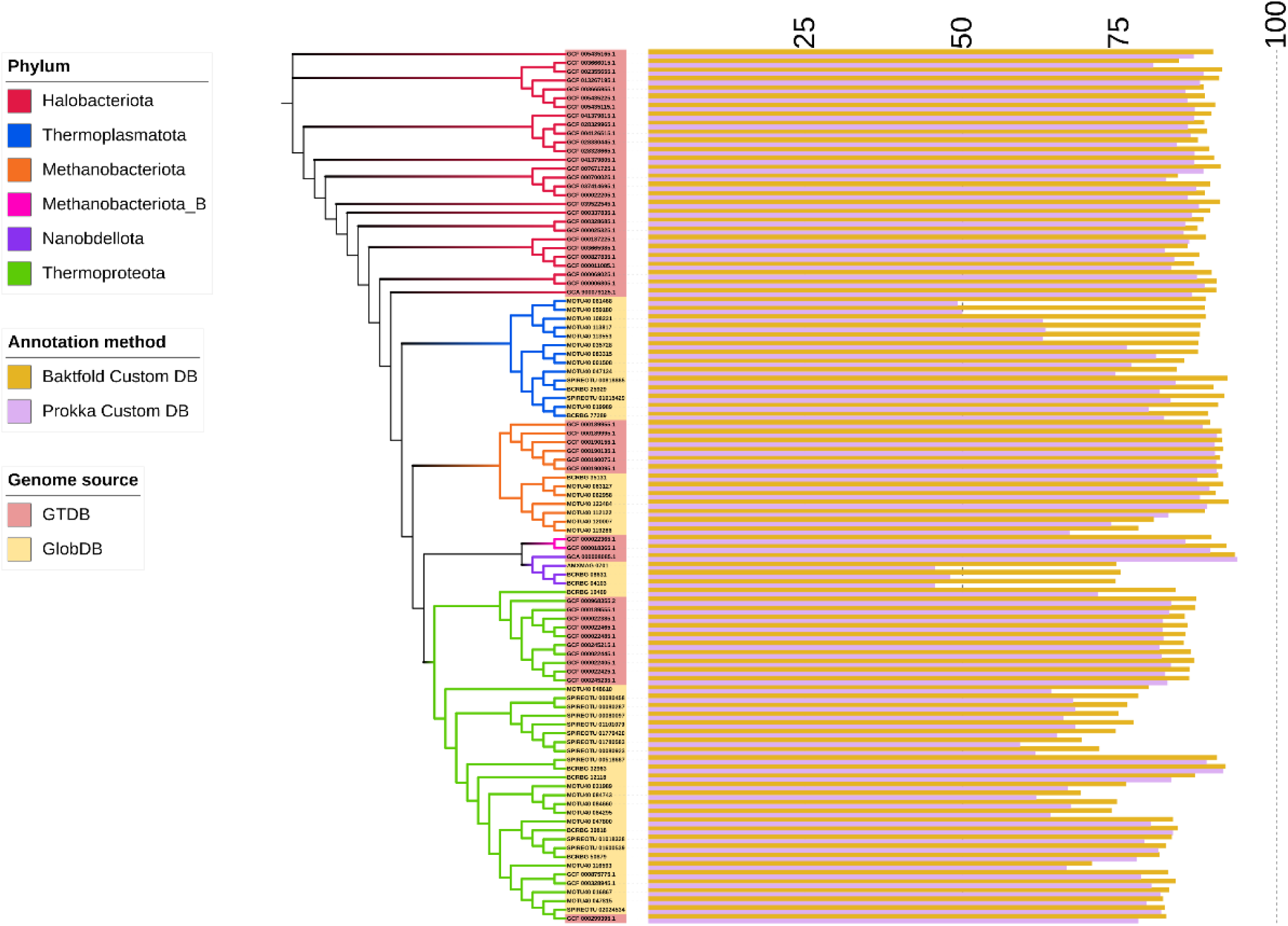
Phylogenetic trees for 100 curated archaeal genomes from GTDB and GlobDB (left) as presented in Figure 5 with the functional annotational annotation rate presented as bars to the right of each genome for Baktfold and Prokka, both with our customised archaeal protein database consisting of nearly 2M archaeal proteins.

**Supplementary Figure 5:**
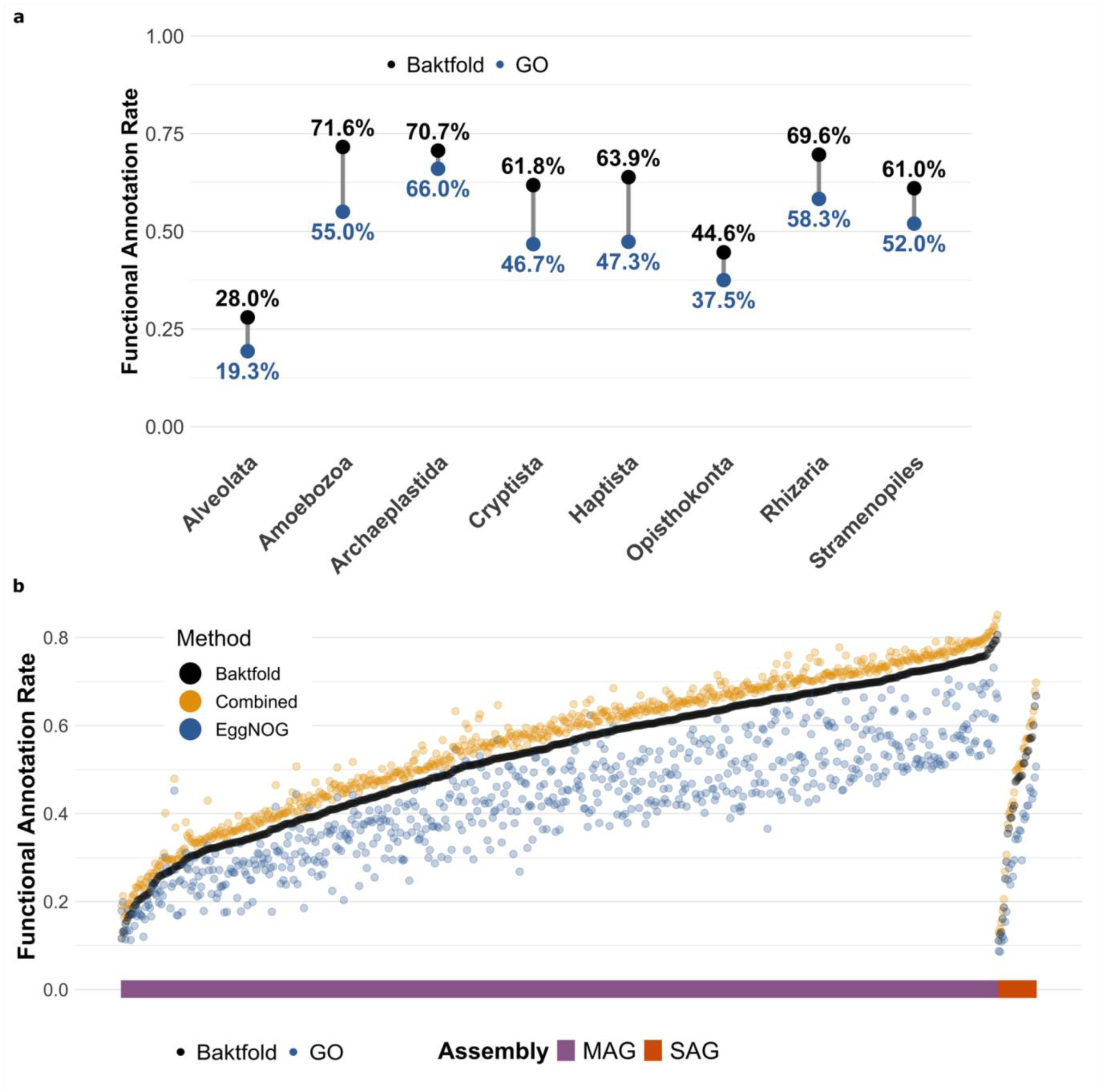
(a) Functional annotation rates of Baktfold (black) and eggNOG-mapper (blue) across 6 taxonomic super groups from the SMAG dataset (b) Functional annotation rates of Baktfold (black) and eggNOG-mapper (blue) and combined (gold) calculated across each of the 713 genomes in the dataset, split by assembly type (MAG or SAG), sorted in ascending Baktfold annotation rate.

## References

1. Hunt M, Lima L, Anderson D, et al. AllTheBacteria – all bacterial genomes assembled, available, and searchable. bioRxiv.; 2025 10.1101/2024.03.08.584059.

2. Chikhi R, Lemane T, Loll-Krippleber R, et al. Logan: Planetary-Scale Genome Assembly Surveys Life’s Diversity. bioRxiv. 2024.07.30.605881; 2025 10.1101/2024.07.30.605881.

3. Schwengers O, Jelonek L, Dieckmann M A, et al. Bakta: rapid and standardized annotation of bacterial genomes via alignment-free sequence identification. Microbial Genomics. 7:000685; 2021 10.1099/mgen.0.000685.

4. Seemann T. Prokka: rapid prokaryotic genome annotation. Bioinformatics. 30:2068–2069; 2014 10.1093/bioinformatics/btu153.

5. Tanizawa Y, Fujisawa T and Nakamura Y. DFAST: a flexible prokaryotic genome annotation pipeline for faster genome publication. Bioinformatics. 34:1037–1039; 2018 10.1093/bioinformatics/btx713.

6. Altschul S F, Gish W, Miller W, et al. Basic local alignment search tool. Journal of Molecular Biology. 215:403–410; 1990 10.1016/S0022-2836(05)80360-2.

7. Buchfink B, Xie C and Huson D H. Fast and sensitive protein alignment using DIAMOND. Nat Methods. 12:59–60; 2015 10.1038/nmeth.3176.

8. Steinegger M and Söding J. MMseqs2 enables sensitive protein sequence searching for the analysis of massive data sets. Nat Biotechnol. 35:1026–1028; 2017 10.1038/nbt.3988.

9. Finn R D, Clements J and Eddy S R. HMMER web server: interactive sequence similarity searching. Nucleic Acids Research. 39:W29–W37; 2011 10.1093/nar/gkr367.

10. Larralde M and Zeller G. PyHMMER: a Python library binding to HMMER for efficient sequence analysis. Bioinformatics. 39:btad214; 2023 10.1093/bioinformatics/btad214.

11. Remmert M, Biegert A, Hauser A, et al. HHblits: lightning-fast iterative protein sequence searching by HMM-HMM alignment. Nat Methods. 9:173–175; 2012 10.1038/nmeth.1818.

12. Cantalapiedra C P, Hernández-Plaza A, Letunic I, et al. eggNOG-mapper v2: Functional Annotation, Orthology Assignments, and Domain Prediction at the Metagenomic Scale. Mol Biol Evol. 38:5825–5829; 2021 10.1093/molbev/msab293.

13. Grigson S R, Bouras G, Dutilh B E, et al. Computational function prediction of bacteria and phage proteins. Microbiology and Molecular Biology Reviews. 89:e00022–25; 2025.

14. Vanni C, Schechter M S, Acinas S G, et al. Unifying the known and unknown microbial coding sequence space. eLife. 11:e67667; 2022 10.7554/eLife.67667.

15. Karavaeva V and Sousa F L. Navigating the archaeal frontier: insights and projections from bioinformatic pipelines. Frontiers in Microbiology. Volume 15–2024; 2024 10.3389/fmicb.2024.1433224.

16. Makarova K S, Wolf Y I and Koonin E V. Towards functional characterization of archaeal genomic dark matter. Biochem Soc Trans. 47:389–398; 2019 10.1042/BST20180560.

17. Donvil L, Housmans J A J, Peeters E, et al. In silico identification of archaeal DNA-binding proteins. Bioinformatics. 41:btaf169; 2025 10.1093/bioinformatics/btaf169.

18. Lepper J A, Beryl Rappaport H B and Oliverio A M. Twelve species of human parasites make up half of the literature on microbial eukaryotes. bioRxiv. 2025.08.20.671258; 2025 10.1101/2025.08.20.671258.

19. Jumper J, Evans R, Pritzel A, et al. Highly accurate protein structure prediction with AlphaFold. Nature. 596:583–589; 2021 10.1038/s41586-021-03819-2.

20. Illergård K, Ardell D H and Elofsson A. Structure is three to ten times more conserved than sequence—A study of structural response in protein cores. Proteins: Structure, Function, and Bioinformatics. 77:499–508; 2009 10.1002/prot.22458.

21. Bertoni D, Tsenkov M, Magana P, et al. AlphaFold Protein Structure Database 2025: a redesigned interface and updated structural coverage. Nucleic Acids Res. 54:D358–D362; 2026 10.1093/nar/gkaf1226.

22. Varadi M, Bertoni D, Magana P, et al. AlphaFold Protein Structure Database in 2024: providing structure coverage for over 214 million protein sequences. Nucleic Acids Research. 52:D368–D375; 2024 10.1093/nar/gkad1011.

23. Berman H M, Westbrook J, Feng Z, et al. The Protein Data Bank. Nucleic Acids Res. 28:235–242; 2000 10.1093/nar/28.1.235.

24. Kempen M van, Kim S S, Tumescheit C, et al. Fast and accurate protein structure search with Foldseek. Nat Biotechnol. 42:243–246; 2024 10.1038/s41587-023-01773-0.

25. Rost B. Twilight zone of protein sequence alignments. Protein Engineering, Design and Selection. 12:85–94; 1999 10.1093/protein/12.2.85.

26. Lin Z, Akin H, Rao R, et al. Evolutionary-scale prediction of atomic-level protein structure with a language model. Science. 379:1123–1130; 2023 10.1126/science.ade2574.

27. Mirdita M, Schütze K, Moriwaki Y, et al. ColabFold: making protein folding accessible to all. Nat Methods. 19:679–682; 2022 10.1038/s41592-022-01488-1.

28. Heinzinger M, Weissenow K, Sanchez J G, et al. Bilingual language model for protein sequence and structure. NAR Genomics and Bioinformatics. 6:lqae150; 2024 10.1093/nargab/lqae150.

29. Zhang Z, Wayment-Steele H K, Brixi G, et al. Protein language models learn evolutionary statistics of interacting sequence motifs. Proceedings of the National Academy of Sciences. 121:e2406285121; 2024 10.1073/pnas.2406285121.

30. Heinzinger M, Littmann M, Sillitoe I, et al. Contrastive learning on protein embeddings enlightens midnight zone. NAR Genomics and Bioinformatics. 4:lqac043; 2022 10.1093/nargab/lqac043.

31. Littmann M, Heinzinger M, Dallago C, et al. Embeddings from deep learning transfer GO annotations beyond homology. Sci Rep. 11:1160; 2021 10.1038/s41598-020-80786-0.

32. Boulay A, Leprince A, Enault F, et al. Empathi: embedding-based phage protein annotation tool by hierarchical assignment. Nat Commun. 16:9114; 2025 10.1038/s41467-025-64177-5.

33. Martínez-Redondo G I, Perez-Canales F M, Carbonetto B, et al. FANTASIA leverages language models to decode the functional dark proteome across the animal tree of life. Commun Biol. 8:1227; 2025 10.1038/s42003-025-08651-2.

34. Jha N, Kravitz J, West-Roberts J, et al. Gaia: An AI-enabled genomic context–aware platform for protein sequence annotation. Science Advances. 11:eadv5109; 2025 10.1126/sciadv.adv5109.

35. Gligorijević V, Renfrew P D, Kosciolek T, et al. Structure-based protein function prediction using graph convolutional networks. Nat Commun. 12:3168; 2021 10.1038/s41467-021-23303-9.

36. Ashburner M, Ball C A, Blake J A, et al. Gene Ontology: tool for the unification of biology. Nat Genet. 25:25–29; 2000 10.1038/75556.

37. Krishna A, Simon V and Kohli A. DeepFRI Demystified: Interpretability vs. Accuracy in AI Protein Function Prediction. 2025.

38. Barrio-Hernandez I, Yeo J, Jänes J, et al. Clustering predicted structures at the scale of the known protein universe. Nature. 622:637–645; 2023 10.1038/s41586-023-06510-w.

39. Bairoch A and Apweiler R. The SWISS-PROT Protein Sequence Data Bank and Its New Supplement TREMBL. Nucleic Acids Res. 24:21–25; 1996 10.1093/nar/24.1.21.

40. Waman V P, Bordin N, Lau A, et al. CATH v4.4: major expansion of CATH by experimental and predicted structural data. Nucleic Acids Res. 53:D348–D355; 2025 10.1093/nar/gkae1087.

41. Speth D R, Pullen N, Aroney S T N, et al. GlobDB: a comprehensive species-dereplicated microbial genome resource. Bioinformatics Advances. 5:vbaf280; 2025 10.1093/bioadv/vbaf280.

42. Camargo A P, Call L, Roux S, et al. IMG/PR: a database of plasmids from genomes and metagenomes with rich annotations and metadata. Nucleic Acids Res. 52:D164–D173; 2024 10.1093/nar/gkad964.

43. Shen W, Sipos B and Zhao L. SeqKit2: A Swiss army knife for sequence and alignment processing. iMeta. e191; 2024 10.1002/imt2.191.

44. Kallenborn F, Chacon A, Hundt C, et al. GPU-accelerated homology search with MMseqs2. Nat Methods. 22:2024–2027; 2025 10.1038/s41592-025-02819-8.

45. Parks D H, Chuvochina M, Rinke C, et al. GTDB: an ongoing census of bacterial and archaeal diversity through a phylogenetically consistent, rank normalized and complete genome-based taxonomy. Nucleic Acids Res. 50:D785–D794; 2022 10.1093/nar/gkab776.

46. Chklovski A, Parks D H, Woodcroft B J, et al. CheckM2: a rapid, scalable and accurate tool for assessing microbial genome quality using machine learning. Nat Methods. 20:1203–1212; 2023 10.1038/s41592-023-01940-w.

47. Shaw J and Yu Y W. Fast and robust metagenomic sequence comparison through sparse chaining with skani. Nat Methods. 20:1661–1665; 2023 10.1038/s41592-023-02018-3.

48. Lee M D. GToTree: a user-friendly workflow for phylogenomics. Bioinformatics. 35:4162–4164; 2019 10.1093/bioinformatics/btz188.

49. Letunic I and Bork P. Interactive Tree of Life (iTOL) v6: recent updates to the phylogenetic tree display and annotation tool. Nucleic Acids Res. 52:W78–W82; 2024 10.1093/nar/gkae268.

50. Dyer S C, Austine-Orimoloye O, Azov A G, et al. Ensembl 2025. Nucleic Acids Res. 53:D948–D957; 2025 10.1093/nar/gkae1071.

51. Delmont T O, Gaia M, Hinsinger D D, et al. Functional repertoire convergence of distantly related eukaryotic plankton lineages abundant in the sunlit ocean. Cell Genomics. 2:100123; 2022 10.1016/j.xgen.2022.100123.

52. Karsch-Mizrachi I, Arita M, Burdett T, et al. The international nucleotide sequence database collaboration (INSDC): enhancing global participation. Nucleic Acids Res. 53:D62–D66; 2025 10.1093/nar/gkae1058.

53. Bouras G, Nepal R, Houtak G, et al. Pharokka: a fast scalable bacteriophage annotation tool. Bioinformatics. 39:btac776; 2023.

54. Giltner C L, Nguyen Y and Burrows L L. Type IV Pilin Proteins: Versatile Molecular Modules. Microbiology and Molecular Biology Reviews. 76:740–772; 2012 10.1128/mmbr.00035-12.

55. Thomas N A, Bardy S L and Jarrell K F. The archaeal flagellum: a different kind of prokaryotic motility structure. FEMS Microbiol Rev. 25:147–174; 2001 10.1111/j.1574-6976.2001.tb00575.x.

56. Rodrigues-Oliveira T, Belmok A, Vasconcellos D, et al. Archaeal S-Layers: Overview and Current State of the Art. Front. Microbiol. 8; 2017 10.3389/fmicb.2017.02597.

57. Bouras G, Grigson S R, Mirdita M, et al. Protein structure-informed bacteriophage genome annotation with Phold. Nucleic Acids Research. 54:gkaf1448; 2026.

58. Monzon V, Haft D H and Bateman A. Folding the unfoldable: using AlphaFold to explore spurious proteins. Bioinformatics Advances. 2:vbab043; 2022 10.1093/bioadv/vbab043.

